# Transcriptome-wide analysis of roles for tRNA modifications in translational regulation

**DOI:** 10.1101/154096

**Authors:** Hsin-Jung Chou, Elisa Donnar, H. Tobias Gustafsson, Manuel Garber, Oliver J. Rando

## Abstract

Covalent nucleotide modifications in noncoding RNAs such as tRNAs affect a plethora of biological processes, with new functions continuing to be discovered for even well-known tRNA modifications. To systematically compare the functions of a large set of ncRNA modifications in gene regulation, we carried out ribosome profiling and RNA-Seq in budding yeast for 57 nonessential genes involved in tRNA modification. Deletion mutants exhibited a range of translational phenotypes, with modifying enzymes known to modify anticodons, or non-tRNA substrates such as rRNA, exhibiting the most dramatic translational perturbations. Our data build on prior reports documenting translational upregulation of the nutrient-responsive transcription factor Gcn4 in response to numerous tRNA perturbations, and identify many additional translationally-regulated mRNAs throughout the yeast genome. Our data also uncover novel roles for tRNA modifying enzymes in transcriptional regulation of *TY* retroelements, and in rRNA 2’-O-methylation. This dataset should provide a rich resource for discovery of additional links between tRNA modifications and gene regulation.

## INTRODUCTION

In addition to the four common nucleotides present at roughly equal abundance in RNA – A, G, C, and U – it has long been known that covalently-modified nucleotides are present at lower abundance. A classic example of such a modified nucleotide is the 7-methylguanylate cap found at the 5’ end of eukaryotic mRNAs. Although decades of investigation have defined scores of modified nucleotides in a variety of coding and noncoding RNAs, new nucleotide modifications continue to be discovered. Moreover, the functions of many nucleotide modifications remain obscure – while the chemical and structural properties of specific modified nucleotides are often well-understood, the detailed functional and regulatory consequences of specific modifications remain unknown for the majority of RNA modification events.

Nucleotide modifications are particularly common in transfer RNAs (tRNAs), and it is estimated that ~20% of all tRNA nucleotides are covalently modified (Czerwoniec et al., 2009; El Yacoubi et al., 2012; Phizicky and Hopper, 2015). Modified tRNA nucleotides include relatively common species such as 5-methylcytosine (m^5^C) and pseudouridine (Ψ), as well as unusual and complex nucleotides such as the wobble modification 5-methoxycarbonylmethyl-2-thiouridine (mcm^5^s^2^U), which is generated in a multi-step process by the Elongator complex along with a number of additional factors (Esberg et al., 2006; Huang et al., 2005; Huang et al., 2008). At the molecular level, tRNA modifications have been implicated in a wide variety of processes, including stabilization of tRNA secondary structure (Motorin and Helm, 2010), translation initiation (Liu et al., 2016), decoding (Li et al., 1997; Nedialkova and Leidel, 2015; Zinshteyn and Gilbert, 2013), reading frame maintenance (Lecointe et al., 2002), protection of tRNAs from nuclease cleavage/degradation (Alexandrov et al., 2006; Schaefer et al., 2010), and subcellular trafficking of tRNAs (Kramer and Hopper, 2013). At the organismal level, tRNA-modifying enzymes have been implicated in processes ranging from neurodevelopment to meiotic chromosome pairing to abscisic acid signaling to early development (Phizicky and Hopper, 2010). Interestingly, some tRNA modifications can be regulated in response to environmental conditions, as for example thiolation of specific tRNAs in *S. cerevisiae* is responsive to the levels of sulfur-containing amino acids cysteine and methionine in the growth media, with altered tRNA thiolation affecting the translation of relevant biosynthetic proteins (Laxman et al., 2013). Although these and many other examples of tRNA modification biology have been uncovered over decades of study, the functions of many tRNA modifications remain mysterious.

A handful of recent studies have carried out transcriptome-wide analysis of translation, using ribosome profiling (Ingolia et al., 2009) to illuminate the roles for specific tRNA-modifying enzymes in translation and proteostasis (Deng et al., 2015; Nedialkova and Leidel, 2015; Zinshteyn and Gilbert, 2013). Here, we adopt this approach to systematically study the roles for tRNA modifications in translational regulation in *S. cerevisiae*, using ribosome profiling to generate ribosome occupancy maps for 57 yeast deletion strains. As expected, deletion of genes encoding enzymes that modify nucleotides in the tRNA anticodon caused the most dramatic translational phenotypes, while loss of enzymes responsible for more distant tRNA modifications often had few discernible translational phenotypes. Codon-level analysis in many cases recovered the expected stalling or slowing of ribosomes at codons corresponding to relevant modified anticodons, as well as revealing novel codon-level translational perturbations in a number of mutants. Scrutiny of the dramatic translational phenotypes observed in *trm7*Δ cells revealed a potential role for this gene in methylation of rRNA as well as tRNAs. At the level of individual transcripts, mutant effects on translation of specific genes resulted in a variety of downstream outcomes, both expected (altered transcription of amino acid metabolism genes secondary to altered translation of *GCN4* mRNA) and surprising (impaired heterochromatin-mediated gene silencing). Most surprisingly, we find a novel role for the Elongator complex (and other factors involved in generation of mcm^5^s^2^U) in maintaining expression of transcripts associated with *TY1* retrotransposon Long Terminal Repeats (LTRs). Our data illuminate novel aspects of Trm7 and Elongator function, reveal many new examples of translational regulation by uORFs, and provide a rich source of hypotheses for future study.

## RESULTS

### Ribosome footprinting of mutant yeast strains

Budding yeast encode 73 genes currently annotated to play a role in tRNA modification, of which 14 are essential. Here, we set out to characterize translational phenotypes for the remaining 59 nonessential genes. Haploid deletion mutants for 57 genes (two mutants – *pcc1*Δ and *pus6*Δ – failed quality control several times) were obtained freshly by sporulation of heterozygous mutant diploids to confirm the viability of the deletion in question and to minimize the potential for suppressor mutations. Initial studies revealed aneuploidies in a subset of the mutants, the majority of which we subsequently re-derived and confirmed to be euploid. However, for two mutants – *bud32*Δ and *gon7*Δ, both members of the conserved KEOPS complex (Daugeron et al., 2011; Downey et al., 2006; Kisseleva-Romanova et al., 2006; Srinivasan et al., 2011) – we repeatedly obtained haploid strains bearing an additional copy of ChrIX, suggesting that the KEOPS complex may be essential in our strain background. We included these mutants in the final set of 57 mutants despite this aneuploidy, as many of the dramatic ribosome footprinting phenotypes observed in these two mutants were also observed to a lesser extent in the euploid *cgi121*Δ mutant (which exhibits partial but incomplete loss of the t^6^A modification (Thiaville et al., 2015)), suggesting that many of the observed phenotypes accurately reflect KEOPS function. Nonetheless, we urge caution in interpreting results obtained from *bud32*Δ and *gon7*Δ cells, as observed phenotypes may be secondary to second-site mutations.

**Figure 1A** shows these 57 genes, grouped according to their modified nucleotide product. Note that this list includes not only the catalytic subunits of tRNA modifying enzymes, but other factors that affect a given tRNA modification, such as the phosphatase Sit4 that is required for Elongator function in vivo. In addition, it is important to note that many of the encoded proteins are also known to affect nucleotide modifications on other RNA species such as mRNAs or rRNA, or play more pleiotropic roles in cell biology – the multimethylase-activating scaffold protein Trm112 is required for methylation of tRNAs, rRNA, and elongation factors (Liger et al., 2011). When appropriate, we will discuss the potential for non-tRNA targets as the relevant mechanistic basis for translational changes observed below.

**Figure 1.**
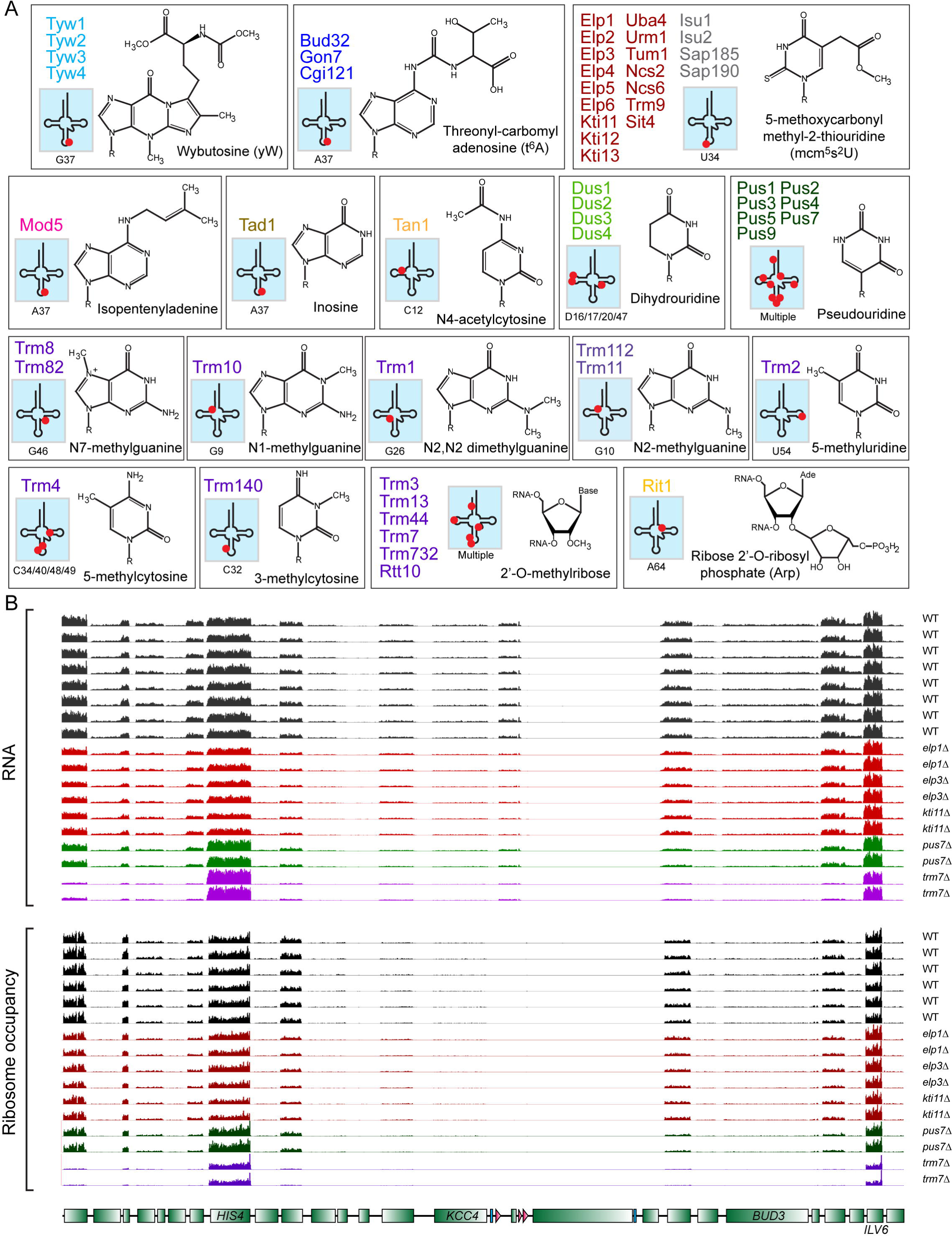
Overview of dataset. (A) Nonessential genes involved in tRNA modifications in budding yeast. Encoded proteins are grouped roughly according to function, as for example the Elongator complex is grouped along with other enzymes required for formation of the mcm5s2U wobble modification. For each group of enzymes, the known product is shown (R indicates ribose in the tRNA backbone for modified bases), along with a tRNA cartoon showing the best-characterized modification locations. For some sets of mutants, the modification shown represents only a subset of products, as for example Elongator and associated factors also catalyze the formation of mcm^5^U, ncm^5^U, and ncm^5^Um, in addition to mcm^5^S^2^U as shown. Modifying enzymes are generally color-coded as indicated here and throughout the manuscript, except in cases where subsets of related factors must be distinguished. (B) Example of RNA-Seq and ribosome footprinting data for chr3:57,000-107,000, showing strong correlations between biological replicate experiments, and also, for the majority of the transcriptome, between mutant strains.

For these 57 deletion strains, we assayed the consequences of loss of tRNA modification on translational control proteome-wide, using ribosome profiling (Ingolia et al., 2009) to provide codon-resolution insight into ribosome occupancy (**Figure 1B**). Matching mRNA abundance data was gathered for each strain as a reference for the ribosome footprinting dataset to enable calculation of translational efficiency per transcript, and to identify mutant effects on transcription or mRNA stability. Wild-type and mutant strains were grown to mid-log phase in rich media and processed for ribosome profiling, with biological duplicates for each strain. **Tables S1-S3** provide complete datasets for mRNA, ribosome occupancy, and translational efficiency, respectively. Overall, biological replicates were well-correlated with one another (**Figure 1B**, **Supplemental Figure S1**), with pairwise correlations ranging from 0.94 to >0.99. Ribosome-protected footprints (RPFs) were predominantly 28-31 nt, as expected, and exhibited known features such as 3 nt periodicity over coding regions and absence of ribosomes over introns. In addition, our data recapitulated prior ribosome profiling analysis of Elongator and other mutants involved in the formation of mcm^5^s^2^U and related modifications (Nedialkova and Leidel, 2015; Zinshteyn and Gilbert, 2013) – see below. This dataset thus provides a high-quality resource for analysis of the roles for tRNA modifying enzymes in translation.

Below, we analyze the dataset at three levels of granularity: averaged across codons, averaged over metagenes, and averaged across individual genes.

### Codon-level analysis of ribosome occupancy changes

Changes in tRNA levels or modifications can affect the dwell time of ribosomes on the relevant codon occupying the A site, and this is readily observed as changes in codon-averaged ribosome occupancy. We therefore analyzed global codon occupancy for all 57 mutants, as previously described (Nedialkova and Leidel, 2015) (**Figure 2A, Supplemental Figure S2A, Table S4**). Our data recapitulate recent studies of mutations affecting mcm^5^s^2^U formation (henceforth collectively referred to as Elongator-related mutants) that documented increased A site ribosome occupancy over AAA, CAA, and GAA codons, which are decoded using mcm^5^s^2^U34-containing tRNAs (Nedialkova and Leidel, 2015; Zinshteyn and Gilbert, 2013), thus validating our dataset (**Figure 2A-B**).

**Figure 2.**
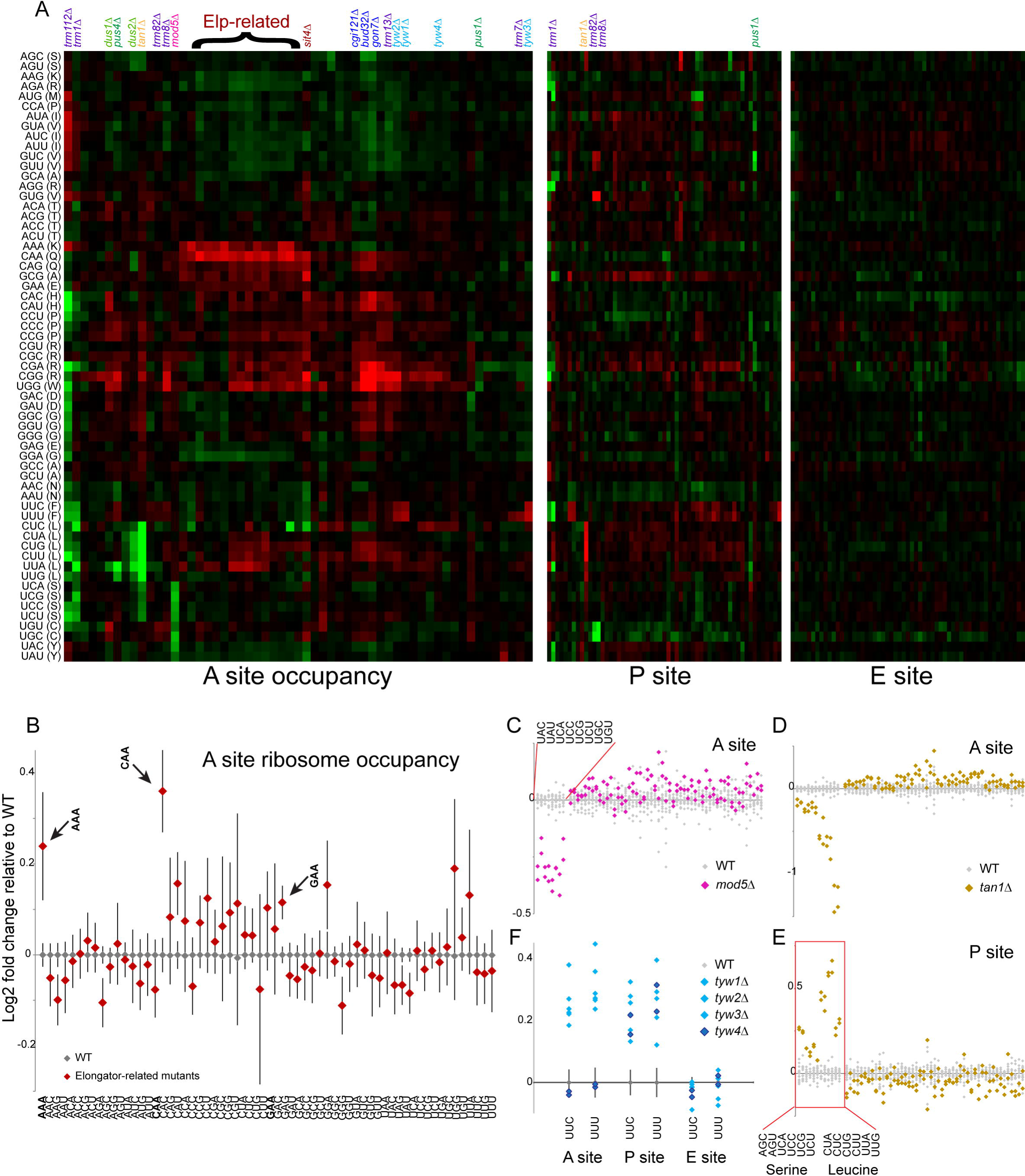
Codon-level analysis of ribosome occupancy. (A) Effects of all 57 mutations on average ribosome occupancy at A, P, and E sites over all 61 codons (excluding stop codons). Columns depict mutations, with key mutations identified above clusters – **Supplemental Figure S2A** shows an expanded view with all 57 mutants annotated, and the entire dataset is available as **Table S4**. Heat maps show log2 fold changes relative to the wild-type average (red=increased codon occupancy; green=decreased codon occupancy). Data for A, P, and E sites are all sorted identically, based on A site dataset clustering, and key mutants are indicated above A and P site clusters. Note that P and E site panels are scaled to 50% of the width of the A site panel. (B) A site ribosome occupancy for all 61 codons for Elongator-related mutants. Grey diamonds show average occupancy and standard deviation for 9 replicates of BY4741, red diamonds show average and standard deviation for 30 Elongator-related datasets (2 biological replicates for 15 deletion mutants). The expected increase in ribosome occupancy is confirmed over AAA, CAA, and GAA, as indicated. (C-E) Data are shown as in panel (B), but here data is shown separately for known target codons (left columns, indicated) with all remaining codons then sorted alphabetically. Data for (C-D) and (E) show A and P site occupancy, as indicated. (F) Discrepant behavior between mutants affecting yW synthesis. A, P, and E site occupancy data shown for Phe codons for the four tyw mutants, as indicated – Tyw1-3 are indistinguishable by design, while the discrepant behavior of Tyw4 is visually emphasized.

For many modifiers with known tRNA substrates, we document altered ribosomal A site occupancy for the relevant codons (**Figure 2C, Supplemental Figures S2B-C, Table S4**), consistent with the tRNA modification in question affecting tRNA stability, charging, or codon recognition. For example, loss of Mod5, which generates N6-isopentenyladenosine (i^6^A) at position 37 in a number of tRNAs (Cys-GCA, Ser-NGA, Tyr-GUA) (Laten et al., 1985), results in dramatically *decreased* A site ribosome occupancy over all relevant codons (**Figure 2C**). For the 2’-O-methylase Trm7, we find increased A site occupancy over codons that exhibit wobble base-pairing with target tRNAs (UGG, UUA, UUU), but not over codons decoded via Watson-Crick base pairing (**Supplemental Figure 2B**). In the case of Tan1, responsible for generation of N4-acetylcytidine (ac^4^C) at position 12 of leucine and serine tRNAs (Johansson and Bystrom, 2004), deletion mutants exhibit dramatically decreased A site ribosome occupancy, but a corresponding increase in P site occupancy over the relevant codons (**Figures 2D-E**). The observation suggests that this modification could potentially play some role(s) in peptidyl-tRNA positioning, enhancing translocation of codons from the P to the E site, or delaying translocation from the A site to the P site. Curiously, loss of the dehydroyuridine synthase Dus2, which is responsible for dU20 formation in the majority of tRNAs (Xing et al., 2004), caused a similar (albeit less dramatic) reduction in A site occupancy specifically of serine and leucine codons (**Figure 2A**), although it did not cause the same compensatory increase in P site occupancy. In addition to relatively specific changes occurring over known target codons, we observed more widespread changes in A site occupancy in mutants lacking Trm1 or Trm112 (**Figure 2A**), consistent with the broad substrate range for these methylases – Trm1 generates N2,N2-dimethylguanosine (m^2^_2_G26) in the majority of cytoplasmic tRNAs (Ellis et al., 1986), while the Trm112 methylase scaffold is required for appropriate methylation of tRNAs, rRNAs, and translation factors (Liger et al., 2011). Interestingly, in both of these mutants, A site occupancy tends to increase over codons beginning with purines, and decrease over codons beginning with pyrimidines (**Figure 2A**).

In addition to these and other cases (**Table S4**) that confirm and extend expected aspects of tRNA modification, we uncovered a number of surprising changes in ribosome occupancy that suggest novel roles for tRNA modifications in translation (**Figure 2F, Supplemental Figures S2C-E**). For example, in several analyses (here and below) we observed discrepant phenotypes for the four *tyw* mutants, despite the fact that loss of any of the four encoded enzymes completely eliminates wybutosine (yW modification at position 37 in tRNA-Phe) synthesis in vivo (Noma et al., 2006). Here, we noted that all four deletion mutants exhibited increased ribosomal P site occupancy over relevant codons, but that A site occupancy at these codons was increased only in mutants lacking Tyw1-3 (**Figure 2F**). As the yW precursor “yW-72” (lacking a methyl group on the α-carboxyl group and a methoxycarbonyl group on the α-amino group of yW) accumulates in target tRNAs in *tyw4*Δ mutants (Noma et al., 2006), we speculate that yW-72 may be sufficient for appropriate decoding of tRNA-Phe, but that the full yW modification is required for efficient peptidyl transfer and/or translocation. As another example, although we observe significantly increased P site occupancy over valine codons GUC/GUG/GUU in mutants affecting the Trm8/Trm82 heterodimer (Alexandrov et al., 2006), these mutants also exhibit unexpected decreases in P site occupancy of UGC and UGU, which are decoded by tRNA-Cys-GCA (**Supplemental Figure S2D-E**).

Taken together, these analyses recapitulate previously-reported translational deficits and thereby validate the quality of our dataset, as well as illuminating potentially novel functions or targets of various tRNA modifications, providing hypotheses for mechanistic followup.

### Trm7 methylates both tRNAs and rRNAs

Turning from the relatively subtle codon-level ribosome occupancy phenotypes described above to gene-level analysis of ribosome occupancy, we noted a particularly dramatic phenotype in initial surveys of the ribosome footprint landscape. Specifically, yeast lacking the tRNA methylase Trm7 – which is required for 2’-O-methylation at positions 32 and 34 of the anticodon loop of several tRNAs (Pintard et al., 2002) – exhibit dramatic, widespread changes in ribosome occupancy at a large number of 5’ UTRs and 5’ coding regions (**Figure 3A**). This is readily captured in metagene analysis of all yeast ORFs aligned according to start codon, and was unique to *trm7*Δ among all 57 mutations in this study (**Figure 3B**). The increase in ribosome occupancy of 5’ UTRs does not appear to result from decreased specificity in ribosome scanning for start sites, as there was no enrichment in AUG or near-cognate start codons in the subset of 5’ UTRs that exhibited increased ribosome ocupancy in this mutant (not shown). Instead, the observed UTR occupancy is likely to be a consequence of globally reduced translation, which manifests as a significantly decreased abundance of polysomes in *trm7*Δ mutants (**Figure 3C**), consistent with prior reports (Pintard et al., 2002).

**Figure 3.**
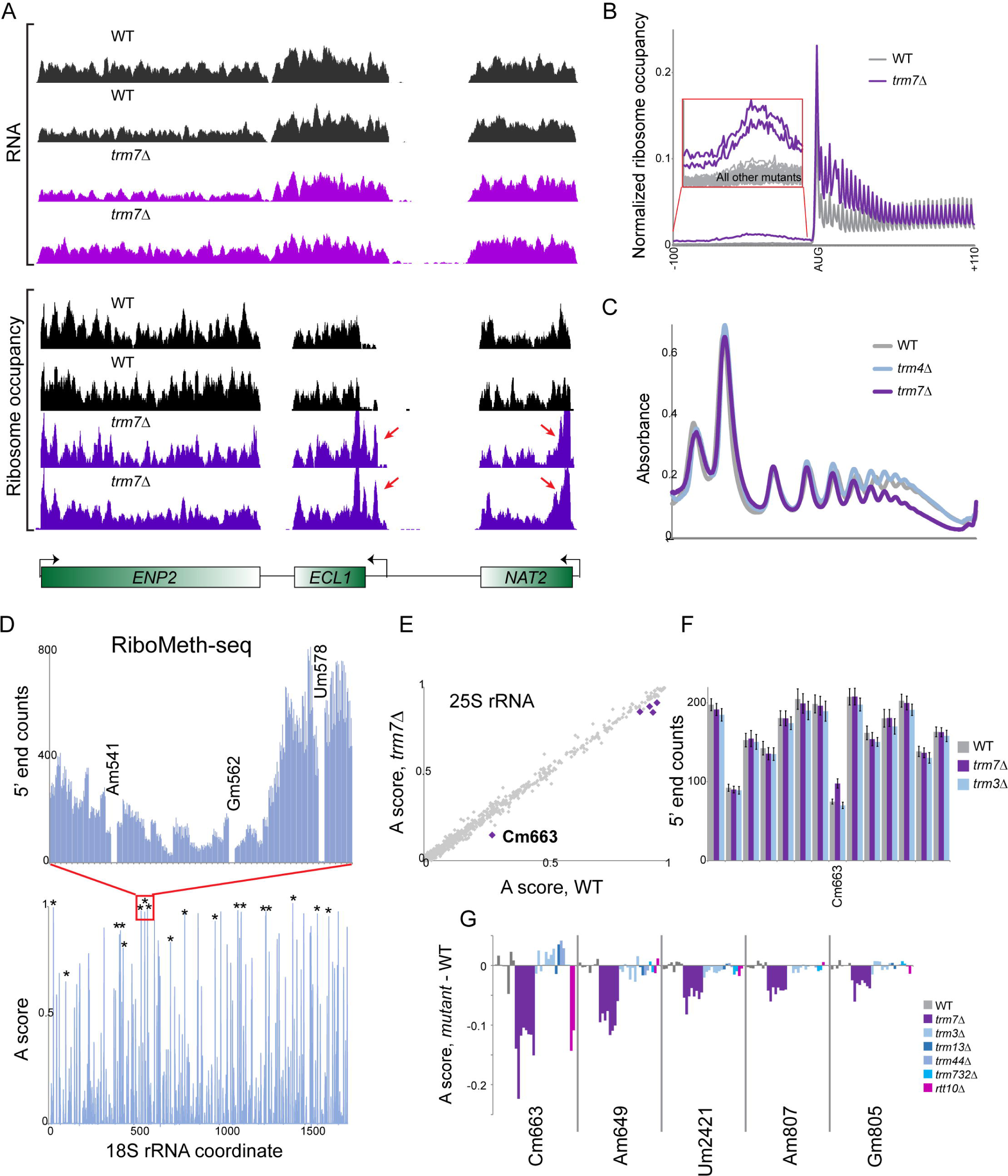
Dramatic effects of Trm7 on ribosome occupancy profiles. (A) Increased ribosome occupancy at 5’ UTRs in *trm7*Δ mutants. RNA-Seq and RPF data for wild-type and *trm7*Δ mutants at characteristic genomic loci. Red arrows show examples of increased ribosome occupancy in mutants. (B) 5’ ribosome accumulation is unique to *trm7*Δ mutants in this dataset. Metagene shows ribosome occupancy data averaged across all genes, aligned by start codon. Main plot shows data for wild-type and *trm7*Δ mutant, while zoom-in on 5’ UTRs shows data for all individual RPF datasets. (C) Polysome profiles of the indicated strains reveal a global deficit in translation in *trm7*Δ. As with the 5’ UTR accumulation in (B), this was specific for *trm7*Δ. WT and unrelated *trm4*Δ are shown for comparison. (D) RiboMeth-Seq analysis of rRNA 2’-O-methylation. Top panel shows counts (normalized to ppm rRNA-mapping reads) of sequencing reads starting across 54 nt of 18S rRNA, as indicated. The three annotated locations are dramatically underrepresented, and correspond to three well-known 2’-O-methylation sites on 18S rRNA. Bottom panel shows methylation “A scores” (Birkedal et al., 2015; Marchand et al., 2016) aggregated for 8 wild-type datasets – individual replicates are nearly indistinguishable – with ^*^ indicating previously-validated methylation sites. Our dataset also recovers known methylation sites on 25S rRNA (**Supplemental Figure S3A**). (E) Trm7 effects on rRNA RiboMeth-Seq. Scatterplot compares methylation A scores for WT (x axis, n=8) and *trm7*Δ (y axis, n=8) strains. The five significantly differentially-methylated nucleotides are indicated with large purple points, and lose methylation in *trm7*Δ but are unaffected in the unrelated *trm3*Δ mutant (**Supplemental Figure S3B**). (F) Normalized RiboMeth-Seq 5’ end read starts (as in (D), top panel) for 14 nt surrounding 25S rRNA C663, as indicated. (G) Comparison of mutant effects on the five candidate Trm7 target sites in 25S rRNA, shown as the change in A score for each mutant replicate relative to the average of 8 WT replicates. For the five Trm*7* target nucleotides, data are shown for WT, trm7, and trm3 mutants (n=8 each), and for t*rm13*, *trm44*, *trm732*, and *rtt10* mutants (n=2 each). Note that C663 methylation is lost in mutants affecting Trm7 as well as one of its heterodimerization partners, Rtt10, while the remaining four potential Trm7 target sites are not affected by either Trm732 or Rtt10.

This dramatic translational phenotype suggested that Trm7 may have additional substrates, particularly since neither of the two known partners of Trm7 – Trm732 and Rtt10 – had similar effects on ribosome occupancy at 5’ UTRs (**Figure 3B**). As the closest Trm7 homolog in bacteria is an rRNA methylase (Pintard et al., 2002), we tested the hypothesis that Trm7 might also methylate rRNA in budding yeast. We assessed 2’- O-methylation of rRNA using RiboMeth-Seq (Marchand et al., 2016), in which limited alkaline hydrolysis of RNA is used to cleave RNAs at all positions with an unmodified 2’ hydroxyl, allowing sequencing-based identification of 2’-O-m ethyl ation sites based on a reduction in sequencing reads starting immediately downstream of the 2’-O-methylated ribose.

Our RiboMeth-Seq data recovered all known 2’-O-methylation sites in yeast 18S and 25S rRNA, with high concordance between 8 replicate libraries (**Figure 3D, Supplemental Figure S3A, Table S5**). Comparing rRNA methylation in WT, *trm7*Δ, and the unrelated *trm3*Δ mutant (n=8 each), we identified five significantly hypomethylated sites, all of which were specific to the *trm7*Δ mutant (**Figure 3E-F, Supplemental Figure S3B-C**). Extending this analysis to several additional mutants, including other 2’- O-methylases as well as Trm7’s known dimerization partners, revealed that four of the five candidate Trm7 target sites were affected exclusively in *trm7*Δ, but also that the strongest candidate – C663 – was in addition hypomethylated in mutants lacking Rtt10, one of the two known heterodimerization partners for Trm7 (**Figure 3G**). Given that the observed depletion of 5’ end reads starting downstream of C663 is not complete (**Figure 3F**), we infer that this nucleotide is likely to be partially-methylated in populations of ribosomes in the cell, perhaps suggesting a role for Trm7/Rtt10 in context-dependent methylation of specific subpopulations of ribosomes. Thus, although the basis for the global effects of Trm7 on translation remain unclear, our data do support the hypothesis that the tRNA methylase Trm7 also methylates a subset of rRNA molecules in vivo.

### Transcriptional consequences of loss of tRNA modifications

We next sought to assess how the loss of specific tRNA modifications affects genomic output at the levels of mRNA abundance and ribosome occupancy (and, by extension, translation efficiency) of individual genes. As expected, changes in mRNA abundance were generally reflected in the ribosome occupancy dataset (**Supplemental Figure S4**). We focus first on mRNA levels, as changes in translational efficiency of key regulators (such as transcription factors) can cause widespread physiological and transcriptional changes that are readily appreciated by RNA-Seq, effectively amplifying the signal for biologically important translational regulation events.

Overall, we find robust mRNA abundance changes in roughly half (27/57) of the mutants analyzed, with minimal or no effects on mRNA abundance observed for the remaining 30 mutants (**Figure 4A**). The vast majority of mutants exhibiting substantial gene expression changes are known to affect tRNA modifications in the anticodon itself or immediately adjacent to the anticodon, including 1) ncm^5^U, mcm^5^U and mcm^5^s^2^U at the wobble position (Elongator and related factors), 2) threonylcarbamoyladenosine (KEOPS), 3) wybutosine (Tyw3), 4) isopentenyladenosine (Mod5), 5) pseudouridine (Pus3/Deg1 and Pus7), and 6) ribose 2’-O-methylation (Trm7). These findings are consistent with the expectation that anticodon modifications should have greater effects on the primary tRNA function – decoding mRNAs – than modifications that occur elsewhere in the tRNA molecule. In addition to the factors involved in anticodon modification, a small number of proteins involved in tRNA modifications distant from the anticodon – such as Rit1, which is required for the 2’-O-ribosylphosphate modification at A64 of initiator tRNA that prevents initiator tRNA from participating in translational elongation (Astrom and Bystrom, 1994) – also exhibited robust gene expression changes. Interestingly, several mutants that robustly affected codon-specific translation, such as *trm82*Δ and *tan1*Δ, had only modest effects on gene expression.

**Figure 4.**
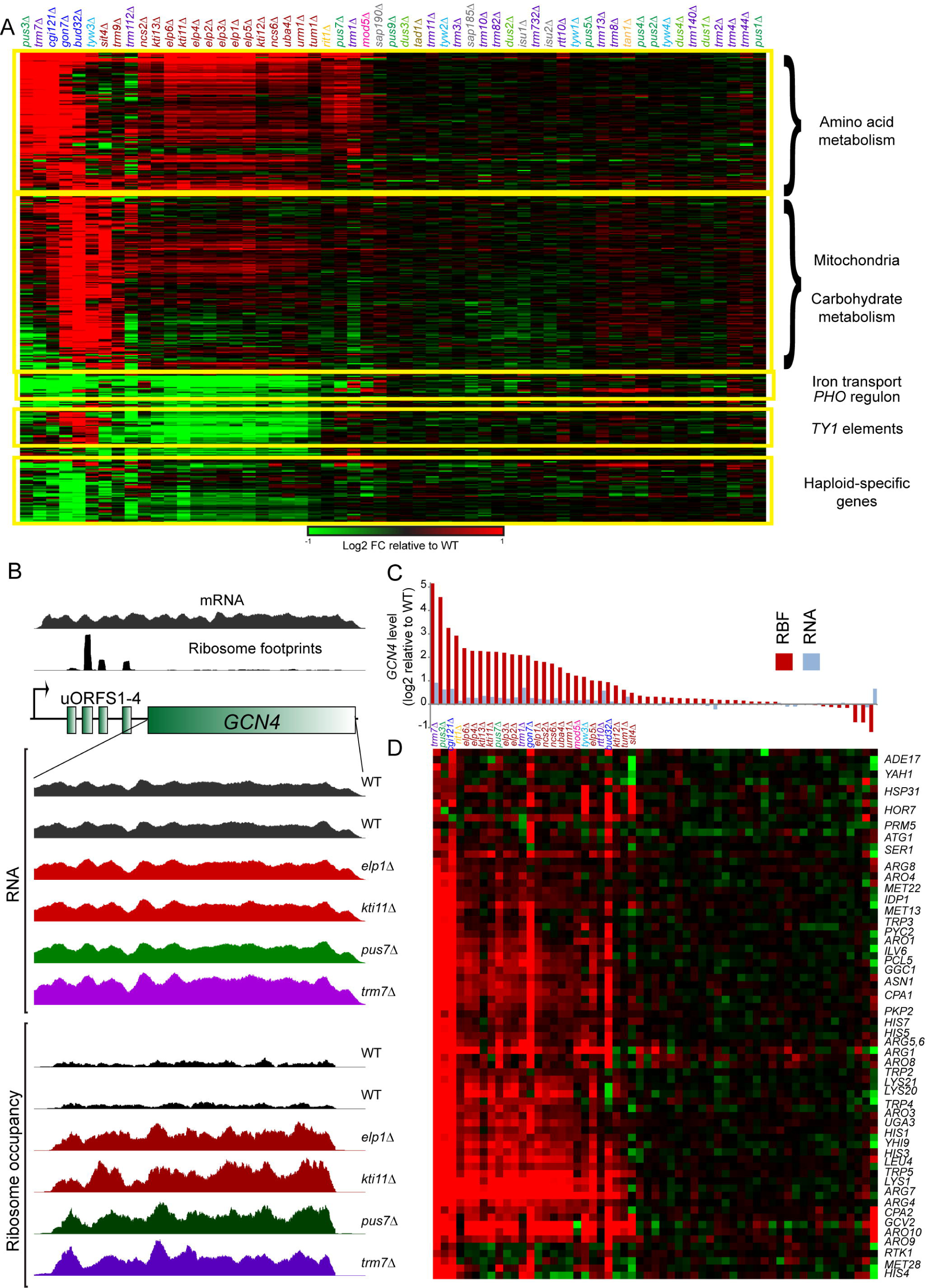
Effects of tRNA modifying enzymes on RNA abundance. (A) Overview of all RNA-Seq changes across the 57 mutants in this study. Data are shown for all genes changing at least 2-fold in at least 2 mutants (filtered for average mRNA abundance > 10 rpkm). Boxes show 5 relatively coherent gene expression clusters, with prominent functional annotations enriched in each geneset indicated. (B) Translational upregulation of *GCN4* is a common occurrence in tRNA modification mutants. Top panels show RNA-Seq and RPF data for the *GCN4* ORF and its 5’ UTR, which carries 4 well-studied regulatory upstream ORFs (uORFs). Zoom-ins show RNASeq and RPF data for indicated mutants, focusing on the *GCN4* coding region. (C) Mutant effects on *GCN4* RNA and RPF levels are shown for all mutants, sorted from high to low Gcn4 translational upregulation. (D) RNA-Seq correlates of GCN4 translational upregulation. Rows show Gcn4 targets (genes that exhibit >2-fold increase in RNA Pol2 occupancy in (Qiu et al., 2016)) for all mutants, sorted as in (C).

Consistent with prior studies on Elongator and Trm9 (Deng et al., 2015; Nedialkova and Leidel, 2015; Zinshteyn and Gilbert, 2013), as well as KEOPS (Daugeron et al., 2011), we observe upregulation of a large group of genes, primarily involved in amino acid biosynthesis and related metabolic pathways, in these mutants. This upregulation can be attributed to translational upregulation of the nutrient- and tRNA-responsive transcription factor Gcn4 (Hinnebusch, 2005) (**Figures 4B-D**). Our data recapitulate these prior findings, further validating the dataset. Moreover, we find that expression of *GCN4* is translationally upregulated in several additional mutants, including *pus3*Δ, *pus7*Δ, *rit1*Δ, *trm1*Δ, *trm7*Δ, *mod5*Δ, and *tyw3*Δ. Thus, a wide variety of aberrations in tRNA function convergently result in increased synthesis of Gcn4, presumably as a consequence of impaired translation of regulatory upstream ORFs (uORFs) in the *GCN4* 5’ UTR. We also find that the upregulation of a proteostasis stress response previously described in Elongator mutants (Nedialkova and Leidel, 2015) is exhibited in additional mutants, with *PRE3* upregulation occurring in mutants affecting Elongator, KEOPS, and Trm112, and *MSN4* upregulation occurring more broadly across the set of mutants that affect the Gcn4 response (**Table S1**).

While the loss of multiple distinct tRNA modifying complexes induced a common transcriptional response through *GCN4* upregulation, other gene expression changes were confined to a more limited set of mutants (**Figure 4A**) and thus were clearly not secondary to increased cellular levels of Gcn4. Among upregulated genes, a large group of genes related to mitochondrial function and carbohydrate metabolism were upregulated in KEOPS mutants and in *tyw3*Δ and *sit4*Δ mutants. Importantly, this was not a result of these strains having lost their mitochondrial DNA, as we detected abundant transcripts for mitochondrially-encoded genes in all of these strains. Other potential explanations for the physiology underlying this gene expression program include altered mitochondrial function resulting from loss of mitochondrial tRNA modifications, or altered expression of the respiration-regulating Hap4 transcription factor. However, in several mutants it is unclear whether this gene expression signature results from loss of the relevant tRNA modifications. Most notably, although the cell cycle phosphatase Sit4 is required for formation of mcm^5^s^2^U (Huang et al., 2008), *sit4*Δ is the only Elongator-related mutant exhibiting the carbohydrate/mitochondria transcriptional phenotype. Similarly, although all four Tyw proteins are required for wybutosine formation, only *tyw3*Δ mutants (which accumulate tRNAs modified with the “yW-86” intermediate (Noma et al., 2006)) upregulate *HAP4* and related genes. On the other hand, this phenotype might potentially reflect a bona fide consequence of loss of t^6^A in KEOPS-related mutants – although the aneuploid *bud32*Δ and *gon7*Δ strains exhibit much stronger upregulation of carbohydrate metabolism genes than do *cgi121*Δ mutants, *cgi121*Δ mutants, which exhibit 80% of wild-type levels of t^6^A, do exhibit a modest effect on these genes (**Figure 4A**). Given the unusual pattern of mutants exhibiting upregulation of carbohydrate metabolism genes, we did not further pursue this connection, although it may prove (at least in the case of KEOPS) an interesting area for future study.

### Silencing-related phenotypes exhibited by tRNA modification mutants

Our next analyses addressed silencing-related phenotypes in the dataset, as tRNA-modifying enzymes have previously been implicated in silencing of subtelomeric and mating type reporters (Chen et al., 2011; Li et al., 2009), and in telomere capping and recombination (Downey et al., 2006; Peng et al., 2015). Defects in various aspects of heterochromatin silencing in yeast result in separable transcriptional responses, which in turn provide robust proxies for function of the relevant silencing pathways. For example, the dramatic downregulation of haploid-specific genes (such as those encoding the pheromone response pathway) observed in *bud32*Δ and *gon7*Δ mutants (**Figure 4A**) is typical of the “pseudo-diploid” state exhibited by budding yeast mutants that fail to repress the silent mating loci (Rusche et al., 2003).

To systematically explore mutant effects on mating locus and subtelomeric silencing (**Figure 5A**), we plotted expression of haploid-specific genes (HSGs) and subtelomeric genes across all mutants in this study (**Figure 5B**, top and middle panels). We also included *PHO* genes as a separate class (bottom panel) – although several *PHO* genes are located near chromosome ends, we noted that both telomere-proximal and -distal *PHO* genes were downregulated in essentially all mutants that affect *GCN4* mRNA translation (**Figure 4A**). Mutants in **Figure 5B** are sorted according to their effects on HSG expression – as is clear in **Figure 4A**, *bud32*Δ and *gon7*Δ exhibit the most dramatic downregulation of HSGs. A small number of additional mutants – *sit4*Δ, *kti11*Δ, *trm7*Δ, *mod5*Δ, and *cgi121*Δ – showed moderate (~1.5-fold changes) downregulation of these genes, with the majority of mutants, including most of the Elongator-related mutants, exhibiting extremely subtle changes in HSG expression. Derepression of subtelomeric genes was similarly restricted primarily to KEOPS mutants. Downregulation of subtelomeric *PHO* genes such as *PHO89* (**Figure 5A-B**) is shown here to emphasize the distinction between the small subset of mutants that specifically affect silencing-related phenotypes, and the larger class of mutants exhibiting relatively nonspecific phenotypes resulting from changes in Gcn4 synthesis.

**Figure 5.**
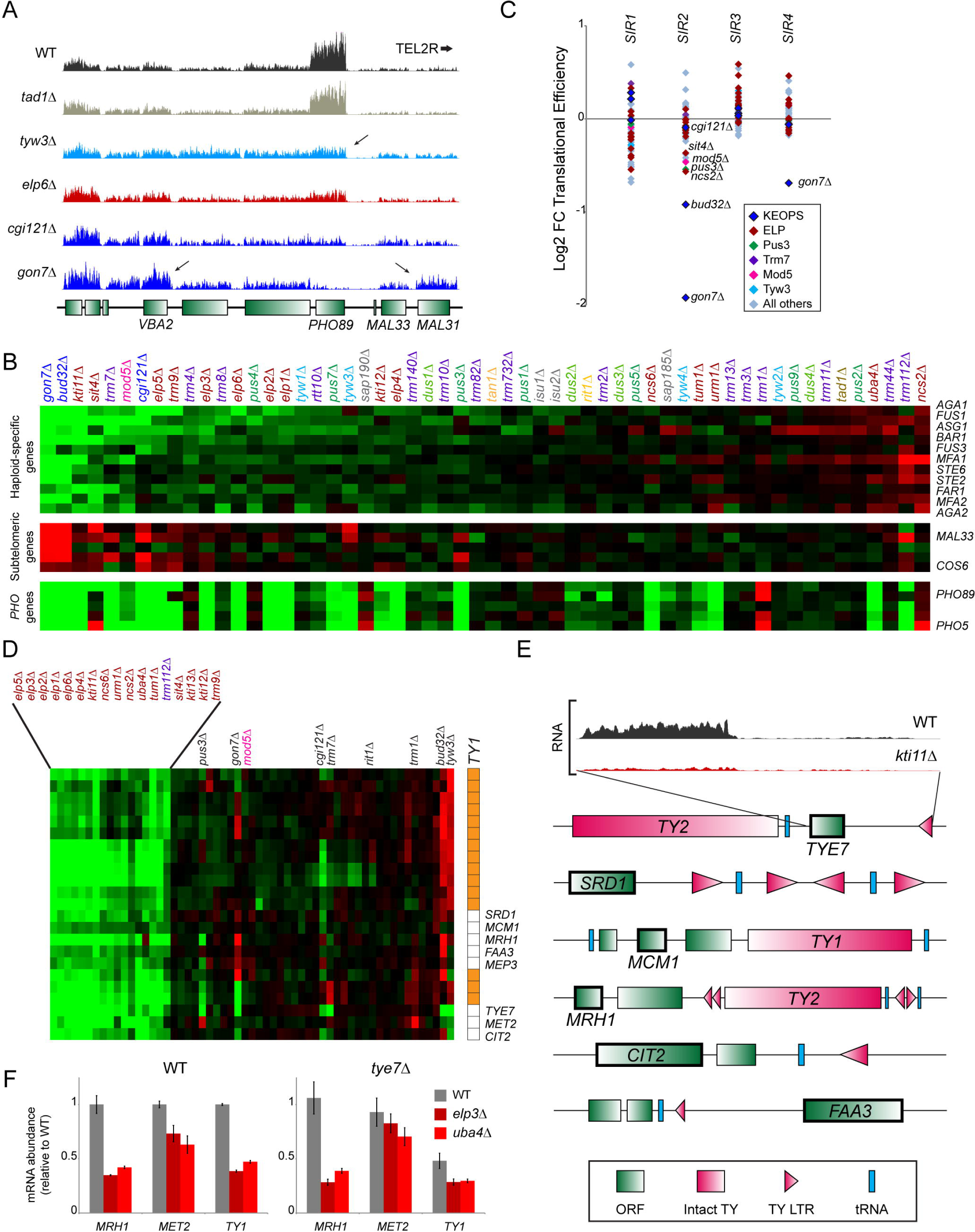
Analysis of silencing-related phenotypes. (A) RNA-Seq data for a ~15 kb locus adjacent to *TEL2R.* Two notable phenotypes are indicated with arrows – repression of PHO genes, observed in a wide range of mutants in this study (**Figure 4A**), and derepression of a subset of subtelomeric genes, which is confined primarily to mutants in the KEOPS complex. (B) Mutant effects on expression of haploid-specific genes (a robust reporter for silent mating locus derepression), subtelomeric genes, and PHO genes, as indicated. Mutants are sorted by their average effect on haploid-specific genes. (C) Mutant effects on translational efficiency of Sir proteins. (D) Downregulation of *TY1* expression in Elongator-related mutants. Cluster shows genes from Cluster 4 (**Figure 4A**) – structural genes encoded by the *TY1* retroelement are indicated with orange boxes. (E) ORFs downregulated in Elongator-related mutants are associated with TY1 long terminal repeats (LTRs). Top panels show RNA-Seq data for TYE7 for WT and a representative Elongator-related mutant. Bottom panels show genomic loci associated with the ORFs shown in panel (D). (F) Elongator effects on target genes are not mediated via changes in TYE7 expression. Q-RT-PCR for two ORFs and for a *TY1* element, as well as two normalization controls *(TEF1* and *TDH3)*, were performed in one of six strain backgrounds – wild-type, *uba4*Δ, *elp3*Δ, *tye7*Δ, *tye7*Δ*uba4*Δ, and *tye7*Δ*elp3*Δ, as indicated – in four replicates. All data are normalized to the wild-type expression levels. Left panel validates our RNA-Seq observations, while right panel shows Elongator effects on these genes in the absence of Tye7. Interestingly, *TY1* mRNA levels are decreased in *tye7*Δ – as expected – but deletion of Elongator leads to a further decrease in *TY1* expression.

As the silencing phenotype previously reported for *elp* mutants was ascribed to defective translation of the *SIR4* mRNA (Chen et al., 2011), we next examined the translational efficiency of the *SIR* mRNAs in our dataset (**Figure 5C**). Consistent with the dramatic silencing defects observed in *bud32*Δ and *gon7*Δ, we found that the translational efficiency of *SIR2* mRNA was significantly decreased in these two mutants (**Figure 5C**). However, outside of this connection, silencing phenotypes were otherwise poorly-correlated with *SIR* mRNA translational efficiency as assayed by ribosome footprinting. For example, although *mod5*Δ mutants did exhibit modest changes in *SIR2* mRNA translation accompanied by moderate downregulation of HSGs, nearly-identical changes in *SIR2* translation in *pus3*Δ and *ncs2*Δ mutants did not result in appreciable mating locus derepression (**Figures 5B-C**).

Taken together, our data indicate that defects in silencing of endogenous loci are relatively rare in mutants that affect tRNA modifications, with the most dramatic silencing defects being confined to the unusual case of the two aneuploid KEOPS mutant strains.

### Regulation of *TY1* expression by Elongator

Although robust silencing defects were largely confined to *bud32*Δ and *gon7*Δ, we uncovered a surprising silencing-related phenotype in Elongator-related mutants. Specifically, we observed substantial downregulation of *TY1* retrotransposon expression occurring almost exclusively in Elongator-related mutants (**Figure 5D**). Interestingly, a small number of endogenous protein-coding genes were also downregulated in the same subset of mutants, and inspection of these genes reveal that all such genes are located in genomic neighborhoods in proximity to intact *TY* elements, solo *TY1* LTRs, and tRNA genes (**Figure 5E**). This link was of great interest to us given the central role for tRNAs in the biology of LTR retrotransposons (Marquet et al., 1995; Weiner and Maizels, 1987), and raise the question of how the Elongator complex affects *TY*-linked gene expression.

Although loss of *TY1* expression occurs, counterintuitively, in *sir* mutants (Lenstra et al., 2011), this is unlikely to explain the downregulation we document for Elongator mutants – KEOPS and other mutants that affect other aspects of Sir-dependent silencing in this dataset do not cause *TY1* repression (**Figure 5D**), and conversely the various Elongator-related mutants exhibit subtle or no effects on other Sir-dependent phenotypes (**Figure 5B**). In addition, the highly Elongator-specific effects on *TY1* regulation cannot be a secondary effect of *GCN4* upregulation, which occurs in a much broader group of mutants (**Figure 4**). Finally, as one of these target genes, *TYE7*, encodes a known transcriptional activator of *TY* LTRs (Lohning and Ciriacy, 1994), we considered the possibility that *TY*-adjacent genes are affected in Elongator mutants as a consequence of altered Tye7 levels. However, q-RT-PCR of several target genes in a *tye7*Δ background revealed further decreases in mRNA abundance in *tye7*Δ*elp3*Δ and *tye7*Δ*uba4*Δ mutants (**Figure 5F**), demonstrating that altered regulation of *TY1* elements is not a secondary effect of Elongator’s effects on the endogenous *TYE7* locus. Given the many links between tRNAs and LTR element replication (Marquet et al., 1995), it will be of great interest in future studies to determine how Elongator functions to support expression of genes located near *TY* LTRs.

### Gene-specific changes in translational efficiency reveal novel regulatory uORFs

We finally turn to analysis of translational efficiency in our dataset. Although mutant effects on the translational control of key regulatory genes, such as *GCN4*, result in an amplified response at the level of mRNA abundance, altered synthesis of many proteins is not expected to cause dramatic transcriptional phenotypes and thus mutant effects on translational efficiency must be addressed directly. **Figure 6A** shows clustered translational efficiency for all 57 mutants, relative to wild-type. As with mutant effects on mRNA abundance, we noted that mutants that affect nucleotide modifications at or adjacent to anticodons exhibited altered translation of many more transcripts than did mutants affecting distal nucleotides. Interestingly, although we identified a handful of relatively specific translational changes in subsets of these mutants, overall we find that the majority of mutants that affect anticodon modifications tended to affect translation of a common group of mRNAs, suggesting that many of the affected genes respond to some aspect of overall translational efficiency (such as, e.g., efficiency of uORF translation), rather than to levels or functionality of individual tRNAs.

**Figure 6.**
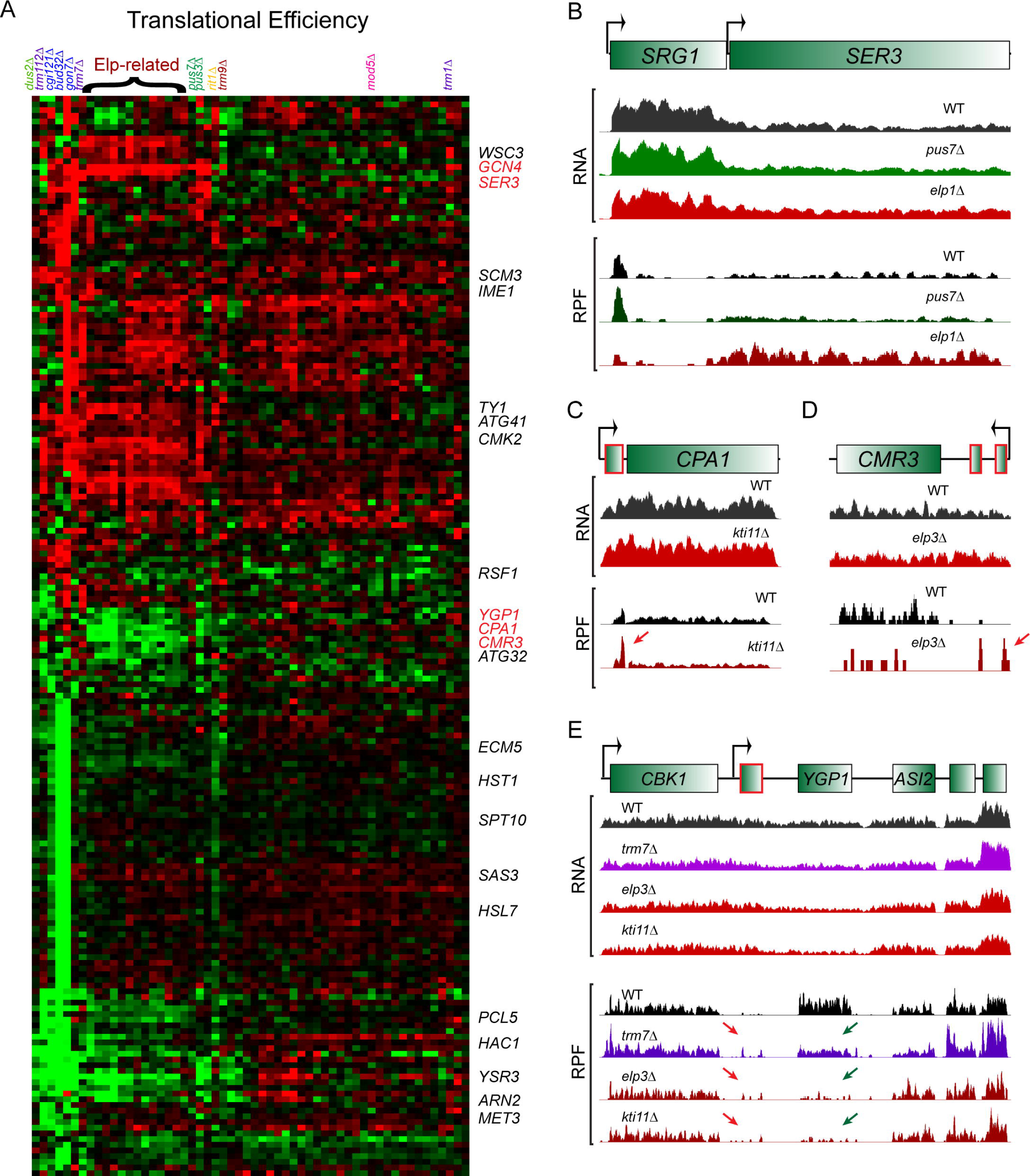
Effects of tRNA-modifying enzymes on translational efficiency. (A) Overview of translational efficiency dataset. Heatmap shows log2 fold changes, relative to wild-type, of all genes with TE changes of at least 2-fold in 2 or more mutants. (B) Translational regulation of *SER3* by uORFs. RNA and RPF (ribosome-protected footprint) data are shown for WT, *pus7*Δ (where SER3 is unaffected) and *elp1*Δ, in which *SER3* translational efficiency is increased. Notable here is a peak of ribosome occupancy over the upstream regulatory transcript *SRG1* which is lost (despite no change in *SRG1* RNA abundance) in mutants that translationally derepress SER3. (C-E) Examples of genes translationally repressed in various tRNA modifying enzyme mutants. Data shown as in panel (B), with green arrows highlighting diminished ribosome occupancy of ORFs, and red arrows highlighting likely regulatory uORFs. Here, known (*CPA1*) or putative (*CMR3*, *YGP1*) upstream regulatory ORFs are highlighted in red in the genomic annotation.

Focusing first on translational ly-u pregulated genes, we noted a small group of genes that exhibited a similar mutant profile to that of *GCN4*. Most notably, we found that translation of *SER3* mRNA was highly correlated with that of *GCN4* across our dataset. Closer inspection of the *SER3* locus revealed clear evidence for ribosome occupancy upstream of the *SER3* start (**Figure 6B**), falling within a previously-described regulatory transcript known as *SRG1* (Martens et al., 2004). Although *SRG1* was originally described as a sense-strand cryptic transcript that is terminated near the *SER3* AUG, Martens *et al.* also noted the presence of long readthrough *SRG1* transcripts extending to the *SER3* 3’ end, and we find multiple sequencing reads spanning the *SRG1*/*SER3* junction, indicating that a subset of *SER3* transcripts include the *SRG1* sequence as their 5’ UTR. These results are most consistent with a model in which translation of *SER3* mRNA is regulated by a uORF in a manner analogous to the intensively-studied mechanism of *GCN4* regulation (Hinnebusch, 2005), and imply that *SRG1* plays separable roles in regulation of *SER3* at both transcriptional and translational levels.

Turning next to translationally down-regulated genes, we noted that *CPA1*, which is known to be translationally regulated by an upstream “attenuator peptide” (Gaba et al., 2005), was downregulated in essentially the same broad set of mutants that affect *GCN4* translation (**Figures 6A, C**). A number of other transcripts were translationally repressed in the same set of mutants that affected *CPA1*, and in many cases we found evidence for uORFs that likely confer translational regulation on the downstream ORFs (**Figures 6A, D, E**). Together, these data provide an expanded survey of presumptive regulatory uORFs, with potential implications for understanding the distinctions between uORFs with stimulatory, vs. repressive, effects on downstream ORF translation.

## DISCUSSION

This dataset provides a unique resource for understanding the roles for tRNA-modifying enzymes and their various cofactors in translational regulation. A key aspect of this dataset is the comparison of multiple disparate mutants within the same dataset, which provides a valuable opportunity to constrain hypotheses for the mechanisms underlying translational phenotypes of interest (see, e.g., **Figure 5**). Overall, we find that those tRNA modifications that occur at or adjacent to the anticodon have the greatest effects on ribosome occupancy, as expected. That said, some modifications that are distant from the anticodon (e.g. position 12), also demonstrate strong effects on ribosome occupancy. The absence of transcriptional or ribosome occupancy phenotypes for many of the remaining mutants involved in tRNA modification could reflect a variety of factors – regulatory feedback could maintain high levels of tRNAs that are destabilized in the absence of a given modification, or a given tRNA modification could play important roles under alternative growth conditions that stress the proteostasis machinery.

More granular analyses at varying levels of resolution from gross transcriptional phenotypes to gene-centric and codon-centric ribosome occupancy reveal both expected behaviors of various mutants as well as novel observations that inform mechanistic hypotheses for future study. We highlight several striking examples of such novel findings, such as the distinction between the *tyw* mutants in A and P site occupancy of relevant codons, and many additional related examples can be found in the various Supplemental Tables.

Most surprisingly, we discover a role for the Elongator complex and other factors required for wobble U modifications (mcm^5^s^2^U and related) in control of expression of both intact *TY1* elements as well as multiple endogenous genes associated with solo LTRs. This finding is of great interest given the ancient links between tRNAs and LTR retroelements – most famously, tRNAs or tRNA-like RNA structures almost universally serve as primers for reverse transcriptase (Marquet et al., 1995; Weiner and Maizels, 1987), and more recent studies implicate cleaved tRNA fragments in control of LTR-associated genes (Martinez et al., 2017; Sharma et al., 2016). We consider a number of hypotheses for the mechanistic basis for Elongator control of *TY1*, ruling out roles for the Sir complex, Gcn4, or Tye7 as mediators of this effect. These findings reveal a novel connection between tRNA modifications and control of LTR elements, and mechanistic dissection of the role for Elongator in *TY* transcription or mRNA stability will be of great interest.

## ACKNOWLEDGEMENTS

We thank A. Korostelev, A. Jacobson, and N. Krietenstein and other members of the Rando lab for discussions and critical reading of the manuscript. This work was funded by NIH grant R01HD080224 to OJR.

## MATERIALS AND METHODS

### Yeast strains and culture conditions

All strains were generated in the BY4741 background (MAT**a** his3Δ1 leu2Δ0 lys2Δ0 ura3Δ0). Haploid deletions were generated by sporulation and tetrad dissection of heterozygous deletions in the diploid BY4743 background, which were obtained either from the Yeast Knockout Heterozygous Collection (Dharmacon), or generated de novo via replacement of genes of interest with KanMX6 in BY4743. Haploid deletions were selected on YPD+G418, Lys minus (SD Glu +His+Leu+Met+Ura), and Met minus (SD Glu +His+Leu+Lys+Ura) media. MAT**a** and deletion genotypes were verified by PCR. After initial analysis of RNA-Seq from all deletion mutants, we identified clear evidence of aneuploidy (consistently elevated expression across entire chromosomes) for several strains. These strains were freshly re-made, and, with the exception of *gon7*Δ and *bud32*Δ for which all isolates obtained carried an addition ChrIX copy, all remade strains were confirmed to be euploid.

### Ribosome profiling

Ribosome profiling was carried out as previously described (Heyer and Moore, 2016; Ingolia et al., 2009), with minor modifications. Briefly, yeast strains were grown to mid-log phase (OD600=0.5-0.6) in YPD (1% Yeast extract, 2% Peptone, 2% Dextrose) at 30°C, treated with cycloheximide for 30 seconds, and harvested by centrifugation. For ribosome footprinting, cells were lysed by glass-bead beating in ice-cold lysis buffer, followed by RNase I treatment. Ribosomes were separated on 10-50% sucrose gradient and 80S monosome fractions were collected and RNA extracted. 27-34 nt ribosome footprints were isolated by denaturing PAGE.

For RNA-Seq, total RNA extracted from the same yeast culture for ribosome footprinting was depleted of rRNA using Ribo-Zero, followed by zinc-based fragmentation for 10 min at 70°C. RNA fragments and ribosome footprints are constructed into deep-sequencing libraries as described in (Heyer et al., 2015). Briefly, RNA 3’ ends were dephosphorylated, ligated with a preadenylated adaptor, and reverse transcribed. cDNA was precipitated and size selected by denaturing PAGE. The purified cDNA was circularized and PCR amplified prior to sequencing.

### Sequencing read mapping and gene-level analysis

Barcoded libraries were pooled and sequenced on an Illumina NextSeq500. Raw fastq reads were de-multiplexed and removed of adaptor sequence using HOMER package (Heinz et al., 2010). RPF reads were mapped to *S. cerevisiae* rDNA and the mapping reads were discarded. The remaining RPF reads and RNA-seq reads were mapped to sacCer3 genome using TopHat v2.0.12 with parameters -p 4 -I 5000 --no-coverage-search. Uniquely mapping reads (filtered by samtools view function and parameter -q 10) in length of 27-34 nt (RPF) or ≥27 nt (RNA-seq) were used for gene-level analysis, in which reads were quantified as raw counts or reads per kilobase of transcript per million mapped reads (RPKM) using HOMER package with open reading frame annotations downloaded from SGD. Translational efficiency (TE) was calculated as relative ribosome footprint density (Ribo-rRPKM) divided by relative RNA abundance (RNA-rRPKM), with rRPKM calculated by normalizing RPKM in each mutant by the average of RPKM in WTs grown in the same batch.

### Codon occupancy analysis

Global codon occupancy analysis was calculated as described in (Nedialkova and Leidel, 2015), with minor modifications. The P-site offset was calculated by examining the cumulative distribution of 28-31 nt reads aligning at start codons. After applying the respective offset to reads of each size, only in-frame reads were used. The first 15 and the last 5 codons of each transcript were removed from the reference. The frequency of each codon in ribosomal A, P, and E sites was calculated for normalization, and divided by the average frequency of the same codon in the three downstream codons from the A site.

### 2’-O-methylation sequencing (RiboMeth-seq)

RiboMeth-seq was carried out essentially as described in (Marchand et al., 2016), except for 3’-end dephosphorylation. For WT, *trm7*Δ, and *trm3*Δ, we initially generated 2 replicate datasets each using T4 PNK, Antarctic phosphatase, or Shrimp alkaline phosphatase for 3’-end dephosphorylation, with no significant effects of any of these variant protocols on methylation. A second round of libraries was built using T4 PNK for 3’ end dephospharylation, with two additional replicates for WT, *trm7*Δ, and *trm3*Δ (final n=4 for T4 PNK protocol, n=8 across all 3 protocols for these three strains), as well as 2 replicates each for *trm13*Δ, *trm44*Δ, *trm732*Δ, and *rtt10*Δ. Methylation A scores were calculated as described in (Birkedal et al., 2015; Marchand et al., 2016).

## SUPPLEMENTAL TABLES

**Table S1. RNA-Seq dataset.**

Sequencing depth-normalized RNA-Seq data for mutant and wild-type (columns), with rows showing the 4884 genes with at least 10 or more reads in all wild-type replicate RPF datasets.

**Table S2. Ribosome footprinting dataset.**

As in **Table S1**, but for ribosome footprinting data.

**Table S3. Translational efficiency.**

Mutant effects on translational efficiency, relative to wild-type, expressed as log2 fold change. Mutants effects on translational efficiency are calculated as RPF rRPKM/RNA rRPKM, with rRPKM calculated by normalizing RPKM in each mutant by the average of RPKM in WTs grown in the same batch.

**Table S4. Codon occupancy.**

Ribosome footprints of 28-31 nt were analyzed as described in **Methods**. Columns show individual replicates of wild type and various mutants, while rows show A-, P-, or E-site occupancy on 61 codons (no stop codons are included in the analysis since the first 15 and the last 5 codons of each transcript were removed from the reference).

**Table S5. RiboMeth-Seq dataset.**

RiboMeth-Seq data for WT yeast and the indicated mutants. Sheets include raw counts of 5’ read starts.

## SUPPLEMENTAL FIGURE LEGENDS

**Figure S1.**
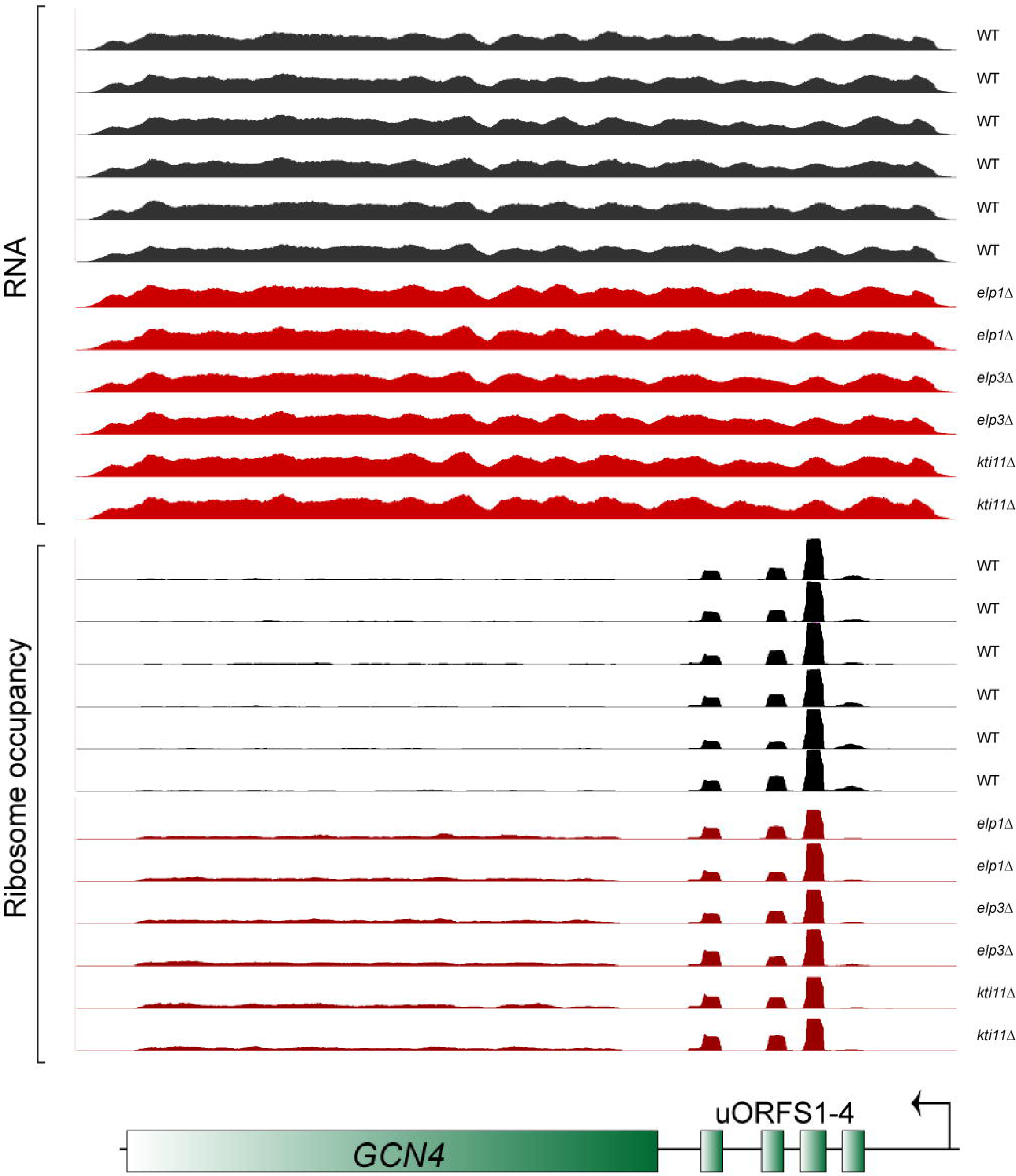
Example of dataset reproducibility. Replicate reproducibility. Data for the *GCN4* locus is shown for the indicated strains, with biological replicates, as well as replicates of distinct but functionally-related mutants, exhibiting highly similar RNA and RBF profiles. Overall all replicate datasets exhibited correlations >0.93, with similarly high correlations observed for datasets for functionally-related mutants (such as *elp1*Δ and *uba4*Δ).

**Figure S2.**
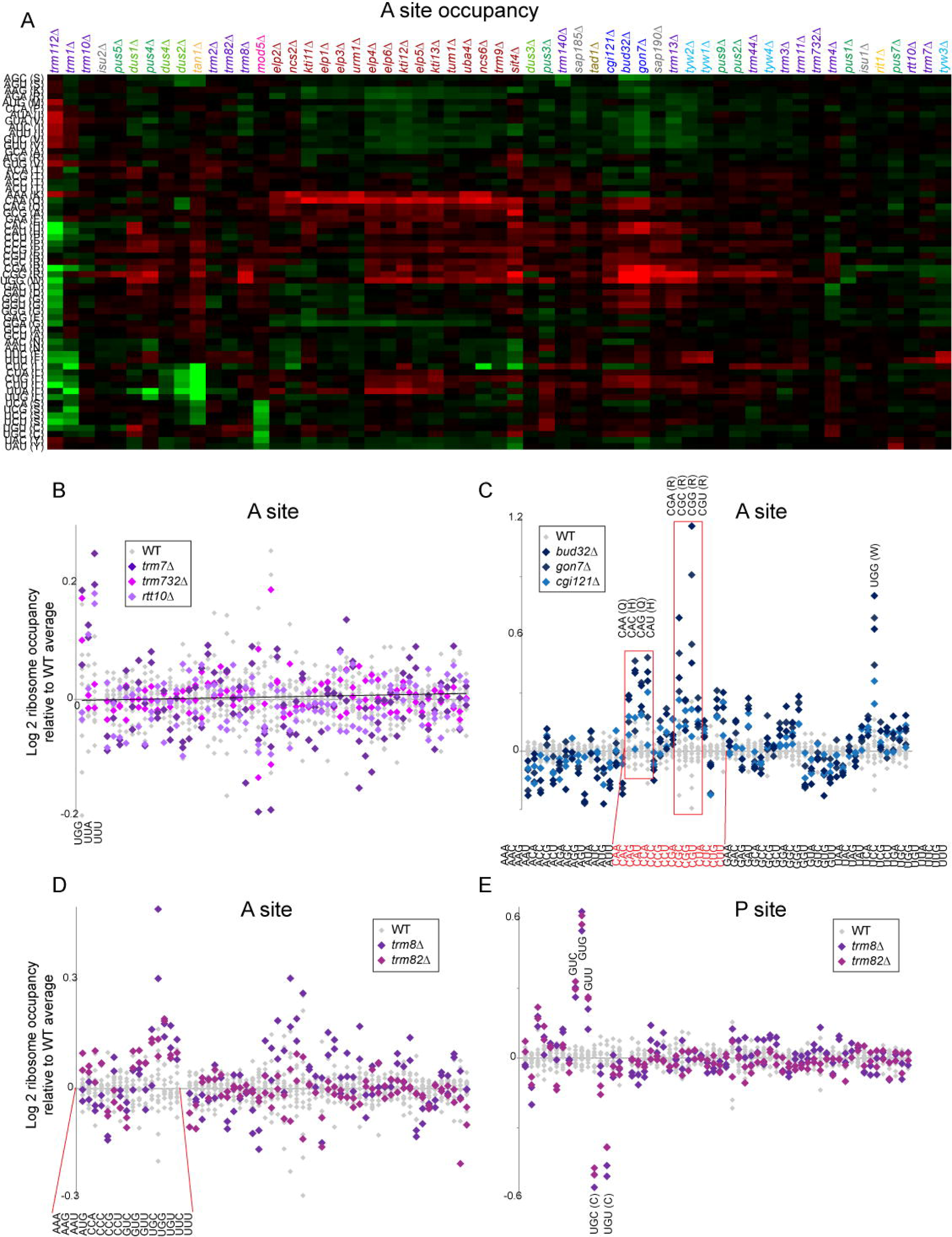
Mutant effects on codon occupancy. (A) A site occupancy panel from **Figure 2A** is reproduced here at larger size to show all 57 mutants. (B-E) Data are shown as in **Figures 2B-E**, with panels (B,D,E) emphasizing relevant codons to the left (with triplet sequences indicated below) and all remaining codons sorted alphabetically, and panel (C) showing all 61 codons sorted alphabetically, as indicated.

**Figure S3.**
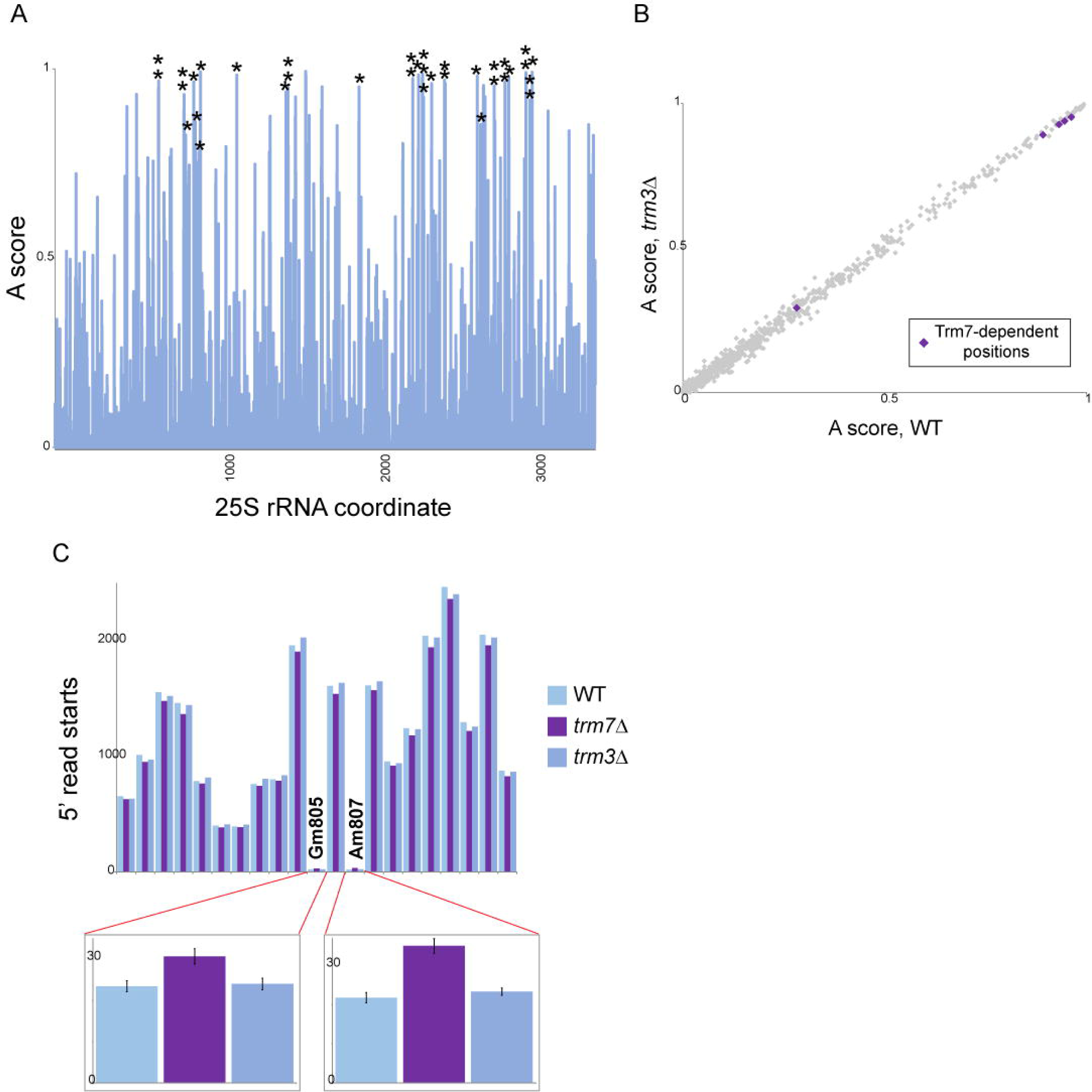
RiboMeth-Seq validation. (A) Validation of 25S dataset. A scores for the WT aggregate dataset are plotted as in **Figure 3D** bottom panel, with previously described 2’-O-methylation sites represented as **\***. (B) Trm7 targets are specific to *trm7*Δ. Scatterplot shows *trm3*Δ effects on 25S methylation scores, with the five Trm7 targets emphasized. (C) Statistically significant effects of Trm7 on highly-methylated rRNA nucleotides represent minor subpopulations of ribosomes. Normalized 5’ end counts shown for a short region encompassing Gm805 and Am807, with insets showing that significant effects of Trm7 here clearly represent an increase in a very small subpopulation of unmethylated ribosomes at these sites.

**Figure S4.**
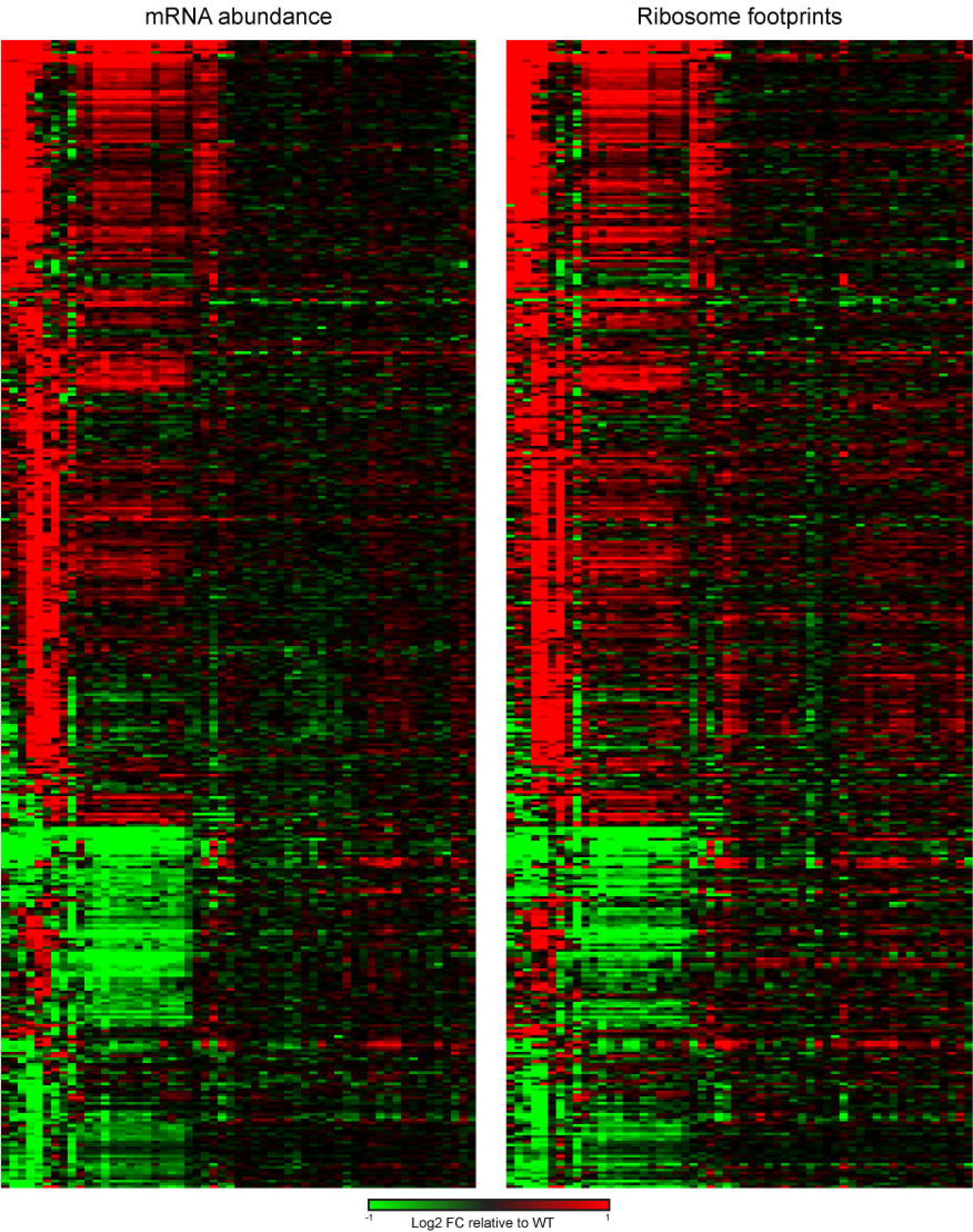
Comparison of RNA-Seq and RPF datasets. RNA-Seq data (left panel) are reproduced from **Figure 4A.** RPF data (right panel) were sorted identically to the RNA-Seq dataset to enable side-by-side comparison.

